# Radiation of pain: Psychophysical evidence for a population coding mechanism in humans

**DOI:** 10.1101/2024.04.02.587666

**Authors:** Wacław M. Adamczyk, Vishwanath Ramu, Catherine Jackson, Geraldine Schulze, Kenneth R. Goldschneider, Susmita Kashikar-Zuck, Christopher D. King, Robert C. Coghill

## Abstract

The spread of pain across body locations remains poorly understood but may provide important insights into the encoding of sensory features of noxious stimuli by populations of neurons. In this psychophysical experiment, we hypothesized that more intense noxious stimuli would lead to spread of pain, but more intense light stimuli would not produce perceptual radiation. Fifty healthy volunteers (27 females, 23 males, ages 14-44) participated in this study wherein noxious stimuli (43, 45, 47 and 49°C) were applied to glabrous (hand) and hairy skin (forearm) skin with 5s and 10s durations. Also, visual stimuli displayed on the target bodily area were utilized as a control. Participants provided pain (and light) spatial extent ratings as well as pain (and light) intensity ratings. In the extent rating procedure, participants adjusted the extent of the square displayed on the screen with the extent of pain (or light) which they experienced. Pain extent ratings showed statistically significant radiation of pain indicated by 12.42× greater spatial spread of pain (pain extent) than the area of the stimulation with 49°C (*p*<0.001), in contrast to visual ratings which closely approximated the size of the stimulus (1.22×). Pain radiation was more pronounced in hairy than glabrous skin (*p*<0.05) and was more pronounced with longer stimulus duration (*p*<0.001). Pain intensity explained only 14% of the pain radiation variability. The relative independence of the pain radiation from pain intensity indicates that distinct components of population coding mechanisms may be involved in the spatial representation of pain versus intensity coding.

## 1. INTRODUCTION

Pain perception is a result of complex neural interactions that take place at multiple levels of the neuroaxis [5]. Unfortunately, our general understanding of how the distributed nociceptive system (DNS) processes nociceptive input and how it contributes to the conscious perception of pain remains incomplete, with the clinical correlate being the fact that many available treatments and prediction strategies have moderate or low effects at best [14,23]. Understanding is particularly limited in spatial aspects of pain where mechanisms of phenomena such as spatial summation of pain [22,34], lateral inhibition [35] and pain radiation [31] remain elusive.

Neural population coding mechanisms have been proposed to encode the intensity of noxious stimulation [5,33,34]. Within the spinal cord the receptive field properties of wide dynamic range neurons (WDR) predict that progressive increases in noxious stimulus intensity would result both in increased discharge frequency of individual neurons as well as a recruitment of activity in progressively larger number of neurons [5,31]. Spinal cord imaging studies confirm this prediction, in both animals [6] and humans [40], such that noxious stimuli recruit more activation than innocuous stimuli and that progressive increases in noxious stimuli result in progressively more spatially extensive activation [5,6,21,40]. During innocuous or mildly noxious stimulation, activity is focused ipsilaterally within a single spinal segment. As noxious stimulus intensity increases, activation is recruited ipsilaterally across multiple spinal segments and also extends to contralateral regions of the spinal cord [5,6].

The role of population recruitment in encoding spatial aspects of pain remains virtually unexplored. Since progressive increases in noxious stimulus intensity recruit progressively more extensive spinal activity, the perceived spatial extent of pain would be predicted to spread as stimulus intensity increases. In one preliminary psychophysical experiment in humans (N=5), noxious heat of graded intensity was applied using an 8mm diameter probe [31]. Pain spread out from the thermal probe despite the fact that surrounding skin temperatures remained unaffected. This radiation of pain in the absence of spread of noxious temperatures was proposed to reflect neural population recruitment as a mechanism involved in the encoding of nociceptive intensity [33], but could also contribute to the perceived spatial dimensions of pain. If population recruitment contributes to spatial aspects of pain, it has major implications for the understanding of chronic pain. Some chronic pain conditions are characterized by pain which is rather well localized, whereas other pain conditions are manifested by widespread radiating pain. For instance, complex regional pain syndrome (CRPS) often starts locally; however, pain can spread extensively over time [36] in contiguous or even non-contiguous manner, with extreme cases of pain spreading to the contralateral limb [19]. Such non-focal distribution of pain [19,34] leads to a mismatch between nociceptive foci and reported symptoms which may confound diagnosis and treatment [42].

To better examine the potential involvement of population recruitment mechanisms in spatial aspects of pain, we aimed to rigorously characterize how pain radiates by measuring pain extent in healthy humans and to test if pain extent patterns are unique (pain versus a visual stimulus) and depend on stimulus intensity, duration and body site (skin type) being tested.

## 2. MATERIAL AND METHODS

### 2.1. Study overview and general information

This study was a mechanistic single-visit investigation performed on healthy individuals. The experiment was based on a within-subject design and involved exposure to noxious heat as well as innocuous visual stimuli (red rectangles). Both modalities of stimuli were delivered to participants’ ventral forearms (hairy skin) and palms of the hand (glabrous skin). The hairy skin site was chosen to be representative of most of the body surface with regards to both primary afferent innervation and spatial acuity. The glabrous skin site was chosen to investigate responses where spatial acuity is the greatest within the somatosensory system [20]. The primary outcome variable was pain extent (“pain size”) rated after each stimulus.

The study procedures were approved by the institutional review board of Cincinnati Children’s Hospital Medical Center (no. 2015-2945). The protocol and analysis plan were preregistered at the “osf.io” platform using AsPredicted.org template (https://osf.io/e8r5f) before data collection started. The study followed principles of the Declaration of Helsinki. After presentation of the objectives, methods, anticipated benefits, and risks, as well as any other relevant aspects of the study, written informed consent was obtained from each participant or legal guardian; participants below 18 provided assent. A monetary remuneration ($100 USD) was given to every participant.

### 2.2. Participants

Participants were recruited via e-mail and social media advertisements as well as word of mouth. They then completed an electronic screener (RedCap) to determine basic eligibility. Then, a clinical coordinator contacted potentially eligible participants and screened them over the phone (or online call) using the inclusion/exclusion checklist. Those participants who were eligible and demonstrated the will to participate were then scheduled for a study visit. To participate subjects had to: i) be healthy, ii) be between 14 and 44 years old, iii) be fluent in written and oral English language, iv) complete study assessments, v) provide written informed consent, and vi) demonstrate willingness to comply with protocol (both parent or guardian and participant). The following exclusion criteria were applied: i) pregnancy, ii) diabetes, iii) history/active chronic pain, iv) psychiatric medications, v) neurological disorders, vi) skin conditions or past skin damage on the arms in or near sites of sensory testing, vii) medications that may alter pain sensitivity or brain activity. Trained research coordinators screened for eligibility. We screened 128 participants and enrolled 54 participants, of which 52 completed experiments **(Appendix)**. Results presented here are from a sample of 50 individuals **(Table 1)** as two participants datasets were not usable due to incomplete subject engagement with the protocol.

**Table 1.** Characteristics of the study sample (N=50)

| Variable | Unit | Mean | SD |
| --- | --- | --- | --- |
| Age | [years] | 24.98 | 7.54 |
| Forearm length | [cm] | 24.78 | 2.10 |
| Hand length | [cm] | 17.93 | 1.35 |
| Fear of pain | [score, 0-100] | 47.20 | 21.60 |
| 2PD at hand | [mm], *H* | 8.93 | 3.53 |
|  | [mm], *V* | 10.94 | 5.55 |
| 2PD at forearm | [mm], *H* | 22.06 | 9.05 |
|  | [mm], *V* | 38.28 | 15.63 |
| Sex | 23 males (46%) , 27 females (54%) |  |  |

Legend: 2PD, two-point discrimination threshold, H, horizontal measurement, V, vertical measurement, SD, standard deviation

### 2.3. Sample size calculations and considerations

For sample size calculation, the project started from the pilot testing on 7 healthy individuals. Participants were attending the experiment based on the same design as described here, i.e., 4 (temperatures) × 2 (stimuli durations) × 2 (body site). The mean and SDs values used for calculation are presented in the **Appendix**. Sample size was then calculated using R and R-Studio (R Core Team, 2023) with implemented “Superpower” library [16]. In brief, this library utilizes the “ANOVA_power” function which generated simulations (here N=1000), based on the means and pooled SD (0.83) from the pilot dataset. Results of this simulation-based power analysis showed that to detect all (significant) main effects in the ANOVA i.e., “temperature”, “duration” as well as “site” with at least 90% power at least 30 participants were needed. However, it was assumed *a priori* that the N = 52 is required to provide a solid reference data for the new outcome, i.e., pain extent measured via the square-matching paradigm (see below).

### 2.4. Equipment and stimuli

Noxious and innocuous heat stimuli were delivered via 16 × 16mm Peltier-based thermode connected to Pathway ATS model (Medoc, Ramat Yishai, Israel). The probe was positioned on the target body region by a 3-degrees of freedom clamping arm (GyroVu, CA, US) attached to a tripod. Heat stimuli were delivered in a ramp up and ramp down format (i.e., trapezoid-shape [34]) with 5s or 10s plateaus. The ramp rates were set to 5°C/s, which allowed for the consistent delivery of thermal stimuli. In addition, an infrared camera suspended over the target region was used to visualize skin temperature externally during the training phase (FLIR One FLIR Systems, Inc., model). Visual stimuli were delivered via a commercially available red LED light externally-voltage controlled via analogue output of digital-analogue converter LabJack U3-HV device (LabJack Corporation, Lakewood, CO, USA). The LabJack converter was controlled by a MacBook Air (Apple Inc., Cupertino, CA) using PsychoPy3 (v2021.2.3) [30] as well as Python3. In general, the visual stimuli had the same temporal and spatial characteristics in the main phase of the experiment, that is, the shape and the extent of the object displayed matched the extent of the thermal probe, and light control had the same ramps and plateau durations as the temperature.

### 2.5. Study structure and design

The experiment was conducted in 4 phases. Following informed consent, participants were first introduced to the rating tasks and were taught in how to operate the trackball (Logitech, Newark, CA, USA) to provide pain extent ratings as well as ratings via computerized Visual Analogue Scale (VAS). They then underwent a well-established familiarization phase [37] in which 32 heat-based stimuli were delivered (temperature range: 35 to 49°C, 5s). Heat stimuli were interleaved by visual stimuli, which also required training (luminance range: 0 to 8 lux ×10^2^).

After familiarization, the second phase assessed 2-point discrimination threshold (2PD) in both tested areas, i.e., hand (glabrous skin) and ventral side of the forearm. The 2PD was determined along the horizontal as well vertical axis using a plastic caliper (Carolina® Biological Supply Company, Burlington, NC). The determination of single 2PD threshold was according to a previously validated protocol [27]. In brief, 4 runs (2 ascending and 2 descending) were performed, however, the first ascending series was based on 5mm steps while the remaining series used on 2mm increments (see [27] for details). Participants reported their sensations using a 2-alternative forced-choice paradigm (1 vs 2 stimuli perceived). The order for the 2PD assessment (body area and stimulus orientation) was counterbalanced and randomly assigned using RedCap software prior to the study visit. Prior to study commencement, research personnel were trained to deliver these stimuli with a pressure that first caused blanching of the skin using approximately a 1s duration of contact.

In the third phase, participants underwent a visual session where visual stimuli of different sizes were presented on the ventral side of the forearm. The aim of this phase was to collect ratings which would allow us to assess the general accuracy of “extent” ratings. In this phase only stimuli with maximal intensity (8 lux ×10^2^) were displayed; however, different masks were used such that the displayed squares had sides of 0.6, 1.6 (as used during the familiarization and the main task), 2.6, or 3.6cm, which corresponded to the displayed areas of 0.36, 2.56, 6.76, 12.96cm^2^. The masks were 3D printed and had a form of filament-based tubes attached to the tip of the light. The masks had square-like holes of dimensions calculated using the law of reversed squares. The masks were always attached in the same distance from the light source and the target (skin) which ensured that the distribution of light was equal across the surface and independent on the size of the cover. Stimuli were applied in a random order and each stimulus size was repeated 3 times to increase measurement accuracy.

The last (main) phase of the experiment was based on 4 blocks of 12 stimuli. In each block, 4 different temperatures were used **(Figure 1)** as a form of heat stimuli (43, 45, 47, 49°C). Each temperature was repeated 3 times, and the order of presentation was pseudorandom. As in the familiarization phase, heat stimuli were interleaved by visual stimuli. Similarly, to heat protocol, 4 intensities of light were applied (2, 4, 6, 8 lux ×10^2^), each of them was repeated 3 times. The selected intensities constituted the possible range of possible brightness output determined by the maximal and minimal voltage input set at the stimulus control device. Thus, in total 12 heat and 12 visual stimuli were delivered per block. Because 2 blocks were applied to the hand (one with 5s and one with 10s plateau), and 2 to the forearm, a total of 48 heat and 48 visual stimuli were used for the main phase of this experiment. Visual stimuli were displayed on the subject’s body instead of an external board to match the attentional demand between sensory systems, i.e. visual and nociceptive.

**Figure 1.**
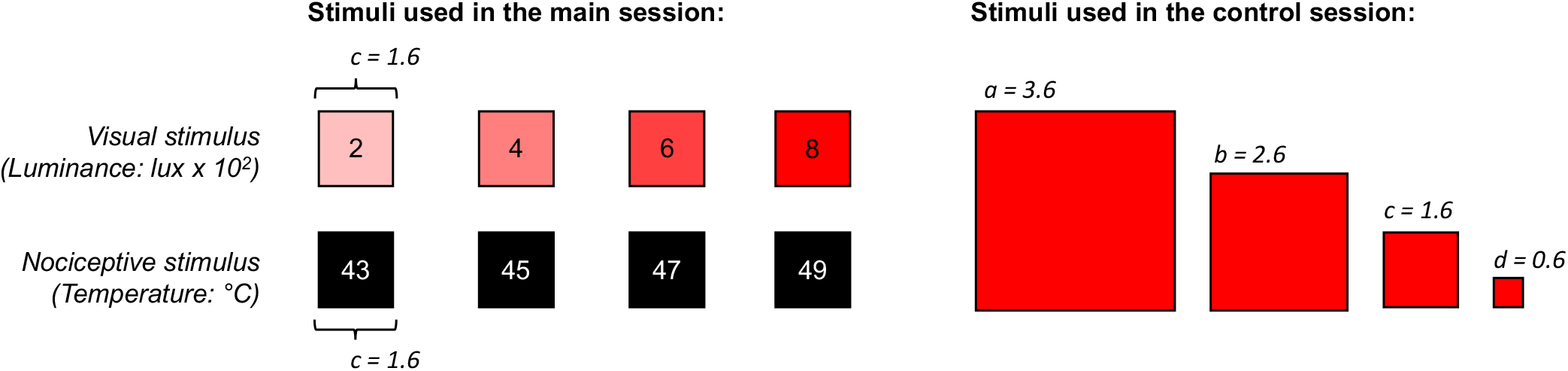
Experimental (heat) and control (light) stimulation. During main phase of the experiment experimental pain was induced using temperature from noxious range (lower left panel: 43 to 49 °C). Each noxious stimulus had the same spatial characteristics (edge “c” = 1.6cm) meaning that the contact with the skin was always the same regardless of the intensity of stimulation. Visual objects of the same size but with different degree of illumination were used as a control (upper left panel: 2 to 8 × 10^2^ Lux). During additional control session visual objects (squares) had different extents (right panel: edge from 0.6 to 3.6cm).

### 2.6. Rating tasks

After termination of the heat stimulus delivery, participants were asked to rate the total pain extent that they felt by adjusting the dimensions of a square displayed on the computer’s screen by moving a trackball. They were told that heat pain can overlap with warmth sensation, but they must only consider the extent of pain, not warmth while rating. After the pain extent rating was provided, the screen was cleared, and then the VAS (**Figure 2**) was displayed. Participants were instructed to first rate pain intensity on VAS with left anchor labelled as “no pain sensation” (0) and right one being “the most intense pain sensation imaginable” (10). The last rating task was with an additional VAS, but they were prompted to rate pain unpleasantness using scale with left anchor set as “no pain sensation” (0) and right one “the most unpleasant pain sensation imaginable” (10). Participants were taught how to distinguish these two dimensions of pain using a sound analogy described elsewhere [32]. Two VAS scales had similar layout and dimensions with differences only related to the anchors and question. Moving the trackball to the right moved a white slider to expose a red bar, mimicking the physical VAS. Thus, the length of the uncovered red bar indicated the magnitude of sensation like in the traditional VAS scale [4]. Python-based code to reproduce task is available on **GitHub**.

**Figure 2.**
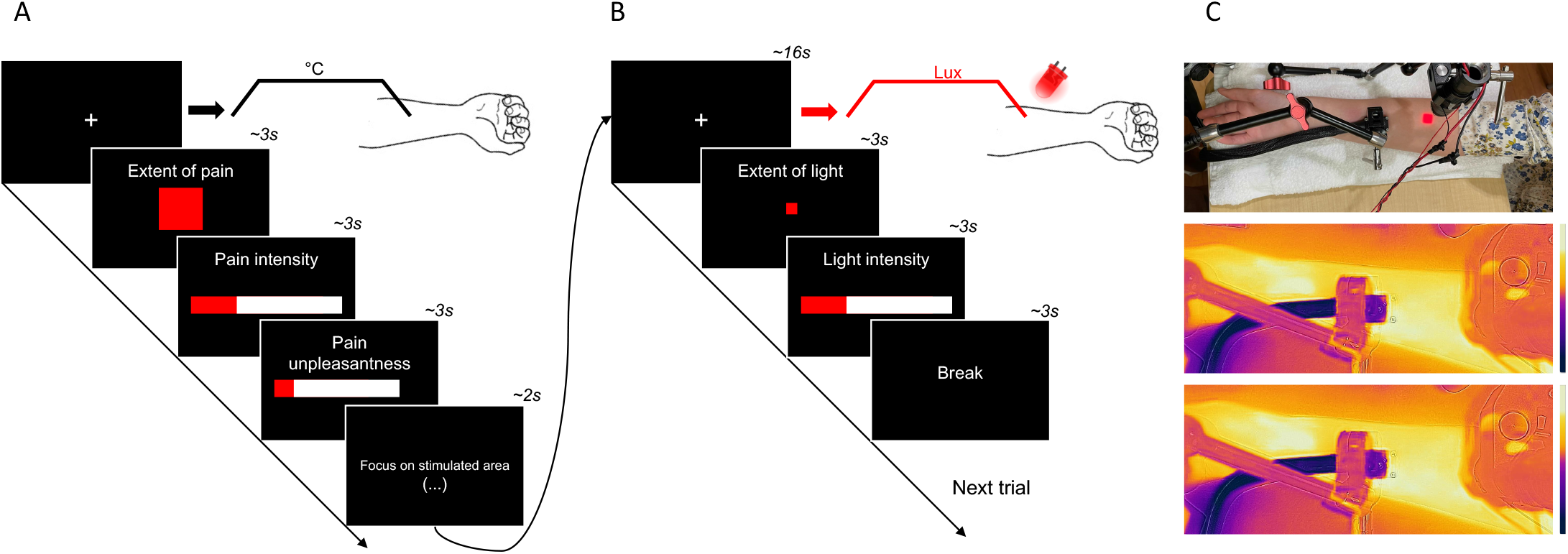
Single trial design. A) Each trial started from a cue presentation followed by heat stimulus (16s, 10s plateau) application to one of the sites (hand vs. forearm). After each stimulus participants rated pain extent (they adjusted the size of square on screen such that it resembled pain extent which they perceived), pain intensity and pain unpleasantness by using computerized Visual Analogue Scale (they adjusted the length of the red bar). B) Subsequently, the 1.6×1.6cm visual stimulus was displayed on their skin which was followed by the light extent and intensity ratings. C) Visual presentation of the stimulus delivery, and skin temperature distribution during 35°C baseline temperature (middle panel) and 49°C (lower panel).

The same process was used after termination of the visual stimulus that was projected onto the participant’s forearm (or hand): the participants rated its perceived extent using the identical trackball-controlled drawing of a red square. Participants were instructed to be as precise as possible, and to evaluate only the extent of light projected onto their skin Participants were also instructed to focus their attention on the light for the entire duration of the light being displayed. Then, they were prompted to rate light intensity on VAS with left anchor set to “no light sensation at all” (0) and right one to “the most intense light sensation imaginable” (10). See **Figure 2** for details.

### 2.7. Data processing and statistical analysis

Collected data were first visually inspected for missing values and quality. No major issues were detected apart from some missing values and erroneous ratings, which were noted during data collection. These typically consisted of erroneous mouse clicks which prematurely terminated the rating procedure before the rating was provided. Some values were also taken out as the patient indicated a lack of attention during the stimulus period which was noted during data collection. Erroneous ratings were documented and set to missing. Average of three (or less) repeated ratings was used in statistical analysis. In the case of missing data, the average of the existing repetitions was used.

The statistical analysis was conducted in the following steps described in the pre-registered protocol. Pain extent ratings were analyzed using a 4 × 2 × 2 repeated-measures Analysis of Variance (ANOVA) with “temperature” (43, 45, 47, 49°C), “site” (forearm, hand) and “duration” (5s, 10s) as within-subject factors. This analysis allowed for testing the primary hypothesis that i) progressive increases in noxious stimulus intensity will elicit greater extent of pain and secondary hypotheses that ii) greater extent of pain will be observed in hairy skin, and iii) when noxious stimuli are longer (10s vs 5s). Main *F* tests were followed by series of post-hoc comparisons (e.g., pain extent in 49°C > in 47°C). Greenhouse-Geisser correction was applied if sphericity assumption was violated, and Bonferroni correction was applied to control type I error rate.

In the second step, the same set of analyses was repeated as in the step 1, but instead of the pain extent ratings, the main dependent variable (DV) was extent of light. This analysis addressed three control hypotheses: it was hypothesized that the i) intensity of visual stimuli will not be related to their perceived extent, ii) the site of body stimulation (hand vs. forearm) as well as iii) the duration of the visual stimuli (5s vs. 10s) will not influence perceived extent of light. Furthermore, to validate the measurement paradigm and to address hypothesis that monotonic increases in the extent of visual stimuli will be related to their perceived extent, one additional ANOVA was performed with “light stimulus” (0.36, 2.56, 6.76, 12.96cm^2^) as a within-subject factor and light extent set as DV.

To validate our results and replicate stimulus-response characteristics of noxious heat [12], analysis from steps 1 and 2 were repeated but with intensity of pain and light set as DVs. Univariate Pearson correlations of pain extent ratings (and light extent ratings) with pain intensity (and perceived light intensity) were performed to address hypothesis that more sensitive individuals will report pain of greater extent. We also tested for potential correlations between pain extent ratings and tactile acuity (2PD thresholds). Exploratory analyses testing for age effects, sex effects and clustering (mixture model) were performed.

Lastly, as a form of psychophysical summary, curve fitting was performed to describe stimulus-response relationship by using Steven’s power function. Individual stimulus-response functions were determined at the subject level and explored through visual assessment for accuracy. Subsequently, the curve was fitted to each individual outcome (intensity, extent, and unpleasantness) within the pain and visual domain (specifically, intensity and extent). The following general power function was initially tested.

***ψ*(*I*) = *k*·*I***^***α***^ where: *I*is a stimulus *k* = intercept, *α* is the exponent, *ψ* is a percept.

The formula was later adjusted towards ‘effective threshold model’ [8] due to poor fitting performance of the original equation [3]:

***ψ*(*I*) = *k*·(*I***−***I***_**0**_)^***α***^ where: *I*_#_ is a first baseline value (e.g., in heat 43°C),

As a result, exponents for each individual were determined by using stimulus data (temperature: 43, 45, 47, 49°C, light intensity in lumens: 2, 4, 6, 8 × 10^2^), and perception data (ratings of participants). Exponents were later extracted at the individual level and averaged across experimental conditions.

Whenever it was possible mean differences were reported with 95% confidence intervals (CI) and effects sizes were reported using Cohen’s *d*_z_ and/or *η*^2^_p_. Pre-processing of the data was performed using MATLAB R2023b (MathWorks, Natick, MA) and all statistical analyses were conducted using IBM Statistical Package for Social Science (SPSS Version 25, Armonk, NY).

## 3. RESULTS

### 3.1. Pain extent ratings

The ANOVA analysis revealed significant effects of “stimulus/temperature” (F_(1.02, 50.16)_ = 14.25, *p* < 0.001, *η*^*2*^_p_ = 0.23, see **Figure 3 and 4)**, indicating that pain extent increased as a function of stimulus intensity and followed the exponential gradient: 43°C vs. 45°C [*p* = 0.10, mean diff. = -0.58, 95% CI: -1.22 to 0.06], 45°C vs. 47°C [*p* = 0.03, mean diff. = -6.27, 95% CI: -12.11 to -0.44], 47°C vs. 49°C [p < 0.01, mean diff. = -18.22, 95% CI: -30.59 to -5.86]. Same analysis revealed also significant main effects of “site” (F_(1, 49)_ = 6.12, *p* < 0.05, *η*^*2*^_p_ = 0.11) and “duration” (F_(1, 49)_ = 12.12, *p* < 0.01, *η*^*2*^_p_ = 0.20) meaning that, in general, i) greater extent of pain was found at the forearm compared to hand and ii) longer stimulus duration led to greater pain extent **(Figure 4)**.

**Figure 3.**
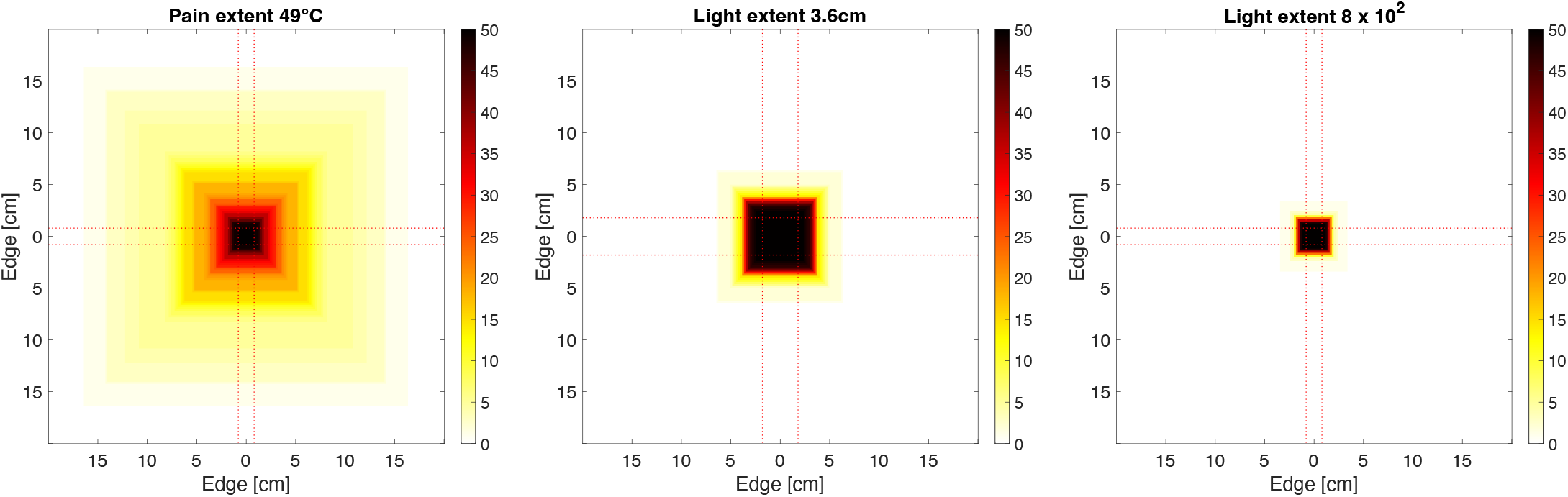
Individual superimposed ratings from selected conditions: >80% of individuals reported different degree of pain radiation (left). Their spatial ratings were very accurate – when light projected the square of 12.96cm^2^ the variability of responses was marginal and overlapped with the actual size of the stimulus (middle). Light perception did not “spread” when the most intense light was used as opposed to pain (left vs. right panel).

**Figure 4.**
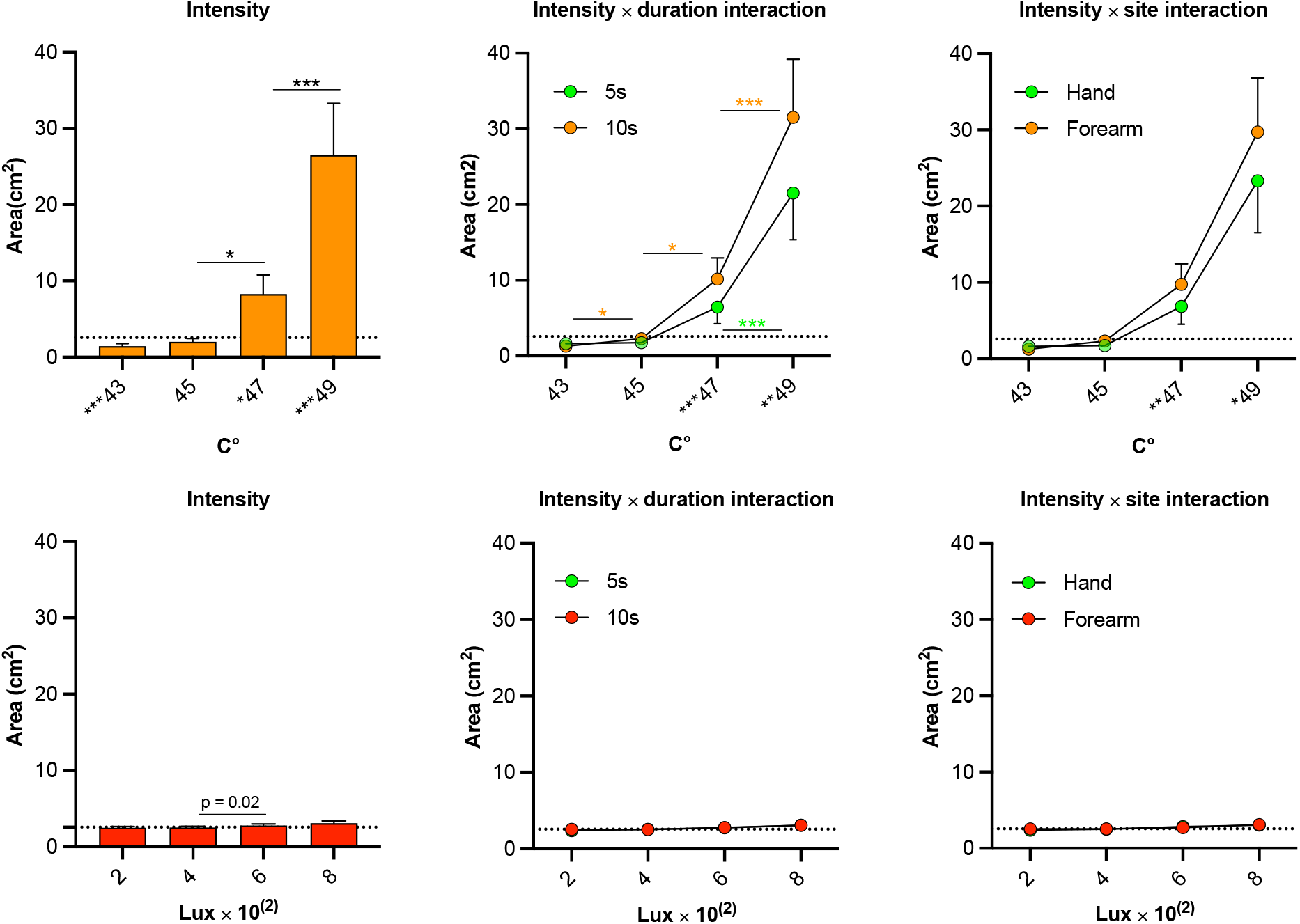
Intensity of heat stimulation (upper panel) and intensity of light stimulation (lower panel) affect extent ratings differently. Pain became more extended as a function of noxious stimulus intensity (upper left) reaching pain extent that was almost 13 times greater than the actual stimulated area (=2.56cm^*2*^) with 49°C. Of note, 43°C*** was related to pain of lesser extent than actual size of the stimulus, whereas 47°C* and 49°C*** stimuli led to pain extent that was greater than the area of stimulation (horizontal line). Significant interaction with stimulus duration (middle) and the site exposed to stimulus (right) indicated that longer stimulus and hairy skin (forearm) produced more pain radiation (greater pain extent ratings) in response to 47 and 49°C but not 43 and 45°C. Light extent (lower panel) ratings remained relatively stable when intensity (luminance) varied (left). No significant interactions with stimulus duration (middle) and site (right) were found. Significance level: ^***^*p* < 0.001, ^**^*p* < 0.01, ^*^*p* < 0.05. Error bars depict standard error of the mean (SEM).

Furthermore, significant “site” × “stimulus” (F_(1.15, 56.14)_ = 3.83, *p* < 0.05, *η*^*2*^_p_ = 0.07) and “duration” × “stimulus” (F_(1.09, 53.22)_ = 10.91, *p* < 0.01, η^2^_p_ = 0.18) interactions were found: Both interactions indicated that i) forearm as stimulated site and ii) duration of noxious heat of 10s led to more pronounced extent of pain independently. Other effects were not statistically significant. Of note, the pain extent was either smaller (43°C: t_(49)_ = -3.41, *p* < 0.01, mean diff. = 1.11, 95% CI: -1.76 to -0.45) or equal to the size of stimulus (45°C: t_(49)_ = -1.22, *p* = 0.23, mean diff. = -0.53, 95% CI: -1.41 to 0.34). However, in the context of the 47° the area at which pain was reported was on average 3.23× larger than actual area (2.56cm^2^) of stimulation (47°C: t_(49)_ = 2.32, *p* < 0.05, mean diff. = 5.73, 95% CI: 0.76 to 10.71), and 10.35× larger when temperature was set to 49°C (t_(49)_ = 3.53, *p* < 0.001, mean diff. = 23.94, 95% CI: 10.30 to 37.57). The pain extent was even 12.42 × larger than stimulation area in the most noxious experimental condition: 49°C delivered for 10s to the forearm (t_(49)_ = 3.97, *p* < 0.001, mean diff. = 31.79, 95% CI: 15.68 to 47.90).

No significant correlations were found between age and pain extent at hand (*r*_*s*_ =-0.04, *p* = 0.81) or forearm (*r*_*s*_ = - 0.08, *p* = 0.57), length of the hand and pain extent (*r*_*s*_ = 0.04, *p* = 0.80) or length of the forearm (*r*_*s*_ = -0.02, *p* =0.88). However, pain intensity was moderately related to the magnitude of the extent of pain intensity explaining less than 25% of the pain extent variability **(see Figure 5)**. Adding unpleasantness to regression model did not affect percentage of the variance explained (49°C, forearm, long pulse: *R*^*2*^ = 0.13 vs *R*^*2*^ = 0.15). No relationship was found between the pain extent (forearm, 10s) and unpleasantness/intensity ratio (*r* = 0.05, *p* = 0.71). No significant sex differences (49°C, forearm, long pulse) were found in pain extent (t_(48)_ = 0.09, *p* = 0.93, *d* =0.03), pain intensity (t_(48)_ = 1.50, *p* = 0.14, *d* =0.43) and pain unpleasantness (t_(48)_ = 1.38, *p* = 0.17, *d* =0.39). Descriptive statistics for these subgroups are provided in the **Appendix**.

**Figure 5.**
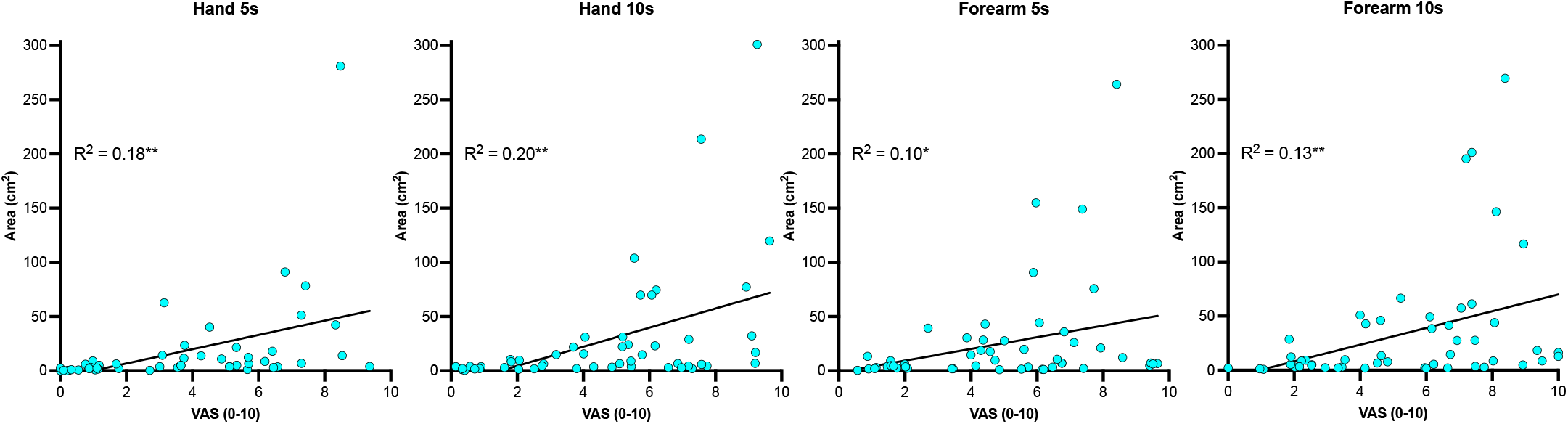
Relationships between pain extent and perceived pain intensity are moderate. Explained variance ranged from 9% (forearm, stimuli of 5s) to 20% (hand, stimuli of 10s). This illustration points that up to 91% of pain extent variability (85% on average) is explained by factors others than percept within the intensity domain. Significance of linear regression line: ^**^*p* < 0.01, ^*^*p* < 0.05.

### 3.3. Intensity of pain

The ANOVA analysis on pain intensity ratings revealed significant effects of “site” (F_(1, 49)_ = 24.58, *p* < 0.001, *η*^*2*^_p_ = 0.33), “duration” (F_(1, 49)_ = 22.70, *p* < 0.001, *η*^*2*^_p_ = 0.32) and “intensity” (F_(1.36, 66.76)_ = 149.59, *p* < 0.001, *η*^*2*^_p_ = 0.75), meaning that, in general, i) more intense pain was found at the forearm compared to hand and ii) longer stimuli duration led to more intense pain. The significant effect of “intensity” indicated gradient of reported pain intensity following the exponential growth: 43°C vs. 45°C [*p* < 0.01, mean diff. = -0.25, 95% CI: -.42 to -0.07], 45°C vs. 47°C [*p* < 0.001, mean diff. = -1.59, 95% CI: -2.04 to -1.14], 47°C vs. 49°C [*p* < 0.001, mean diff. = -2.20, 95% CI: -2.70 to -1.70]. Furthermore, the following significant two-way interactions were found: “site” × “duration” (F_(1, 49)_ = 5.20, *p* < 0.05, *η*^*2*^_p_ = 0.10), “site” × “intensity” (F_(1.99, 97.75)_ = 12.30, *p* < 0.001, *η*^*2*^_p_ = 0.20) and “duration” × “intensity” (F_(2.32, 113.61)_ = 17.61, *p* < 0.001, *η*^*2*^_p_ = 0.26), indicating that i) forearm was more sensitive site over the hand with longer stimulus duration ii) same temperature was perceived differently if it was presented to hand vs forearm iii) and if it was presented using short vs. long stimulation. Lastly, a significant 3-way “site” × “duration” × “intensity” interaction was found (F_(2.47, 120.98)_ = 9.00, *p* < 0.001, *η*^*2*^_p_ = 0.16), meaning that trajectories of increase in pain were modulated by stimulus duration (5s vs. 10s), however this modulation was stronger at the forearm compared to hand (**Figure 6)**. The same pattern of ANOVA results was found in pain unpleasantness **(Appendix)**.

**Figure 6.**
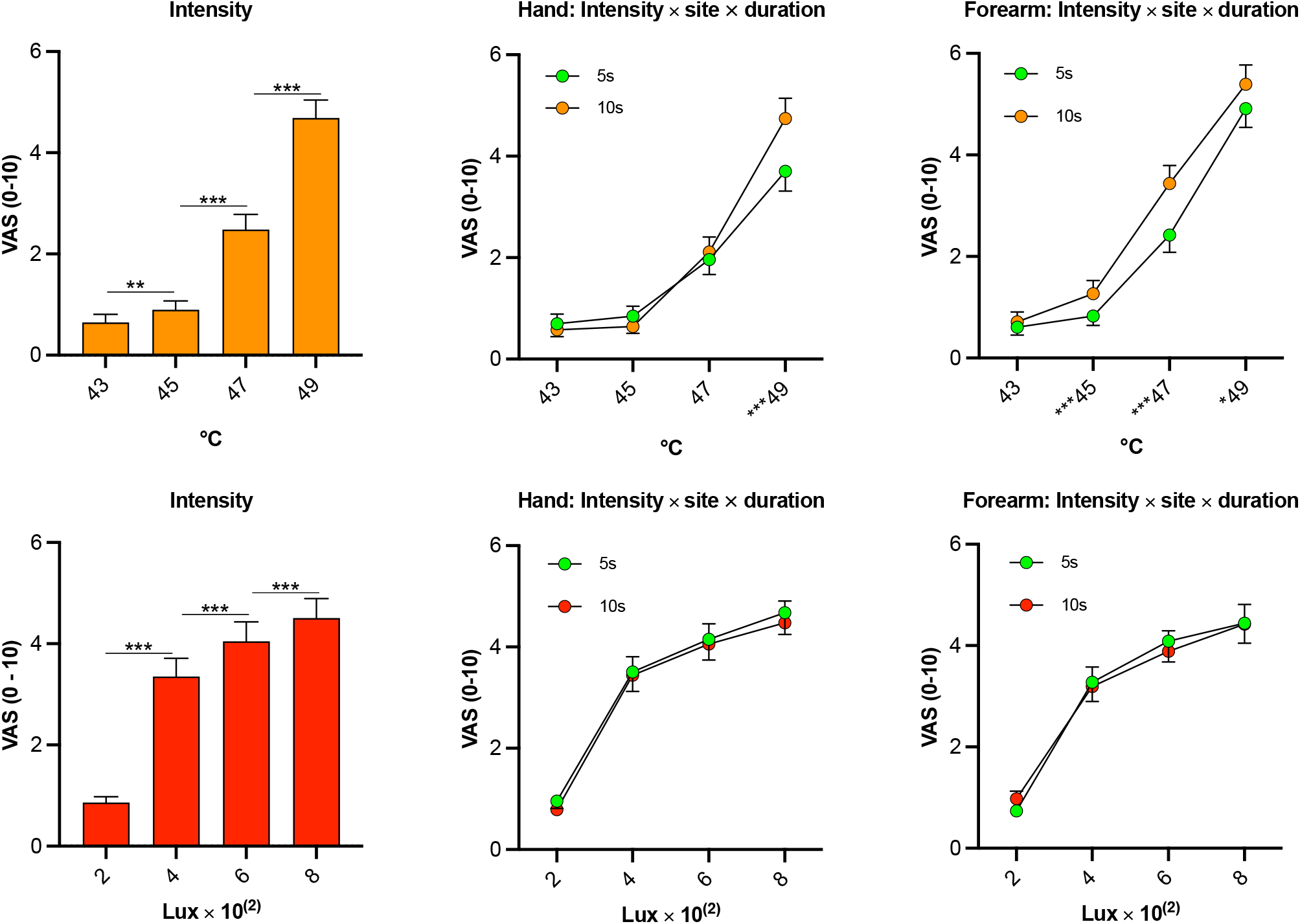
Perceived intensity of pain (upper panel) and light (lower panel) varied as function of stimulus intensity, its duration, site, and their interactions. Pain (upper panel) is perceived as more intense with higher temperatures (left), however, at the forearm (hairy skin) this effect is somehow modulated by duration of the stimulus. Longer duration led to more intense pain induced with 49°C, 47°C as well as 45°C (right). At hand site (glabrous skin) longer duration of stimuli contributed to more intense pain, however, only for 49°C stimulus (middle). Interestingly, perceived light intensity (left) followed logarithmic rather than exponential growth (see Appendix) and was independent on the site and duration of stimulus presentation (middle, right). Three-way interaction was not statistically significant. Significance level: ^***^*p* < 0.001, ^**^*p* < 0.01, ^*^*p* < 0.05. Error bars depict standard error of the mean (SEM).

### 3.3. Perceived light extent

The ANOVA analysis did not show significant effects of “site” (F_(1, 41)_ = 0.19, *p* = 0.66, *η*^*2*^_p_ = 0.004) and “duration” (F_(1, 41)_ = 1.05, *p* = 0.35, *η*^*2*^_p_ = 0.02). However, significant effect of “intensity” was found (F_(1.36, 55.64)_ = 11.57, *p* < 0.001, *η*^*2*^_p_ = 0.22), meaning that, in general, visual stimulus was perceived as more extended when more intense light was displayed on targeted site **(Figure 4)**. This effect, however, seemed marginal when contrasted with pain extent as only one comparison was statistically significant: 2 vs. 4 [*p* = 1.0, mean diff. -0.06, 95%CI: -0.24 to 0.11], 4 vs. 6 [*p* < 0.02, mean difference: - 0.24, 95%CI: -0.46 to -0.03], 6 vs. 8 × 10^2^ lux [*p* = 0.10, mean diff. -0.30, 95% CI: -0.63 to 0.03]. Also, a significant “site” × “duration” interaction was found, likely reflecting minor positioning and technical differences between the light being displayed on different sites at varying distances from the eye (F_(1, 41)_ = 5.59, *p* < 0.05, *η*^2^_p_ = 0.12). No other significant effects were found for this outcome **(Appendix)**. Perceived light intensity was not related to the perceived extent of light within each intensity (*p* > 0.19, r_s_ > 0.19).

### 3.4. Light intensity

The ANOVA analysis on light intensity revealed significant effect of “intensity” (F_(1.47, 60.19)_ = 85.54, *p* < 0.001, *η*^*2*^_p_ = 0.68). Perceived light was reported as more intense with more intense stimulation, as follows: 2 lux vs. 4 lux [p < 0.001, mean difference: -2.48, 95% CI: -3.35 to -1.62], 4 lux vs. 6 lux [p < 0.001, mean difference: -0.70, 95% CI: -0.98 to -0.41], 6 lux vs. 8 lux × 10^2^ [p < 0.001, mean difference: -0.45, 95% CI: -0.75 to -0.15]. No other main or interaction effects were found within light intensity domain (**Appendix**).

### 3.5. Displayed light of different sizes (extents): validation

The ANOVA analysis revealed significant effect of “light size” (F_(1.54, 75.34)_ = 177.86, *p* < 0.001, *η*^*2*^_p_ = 0.78), indicating that participants clearly discriminated different sizes of displayed objects **(Figure 7)**: 0.36cm^2^ vs. 2.56cm^2^ [*p* < 0.001, mean diff. -2.14, 95%CI: -2.68 to -1.59], 2.56cm^2^ vs. 6.76cm^2^ [*p* < 0.001, mean diff. -5.56, 95%CI: -7.24 to -3.88], 6.76cm^2^ vs. 12.96cm^2^ [*p* < 0.001, mean diff. -6.78, 95% CI: -8.40 to -5.15].

**Figure 7.**
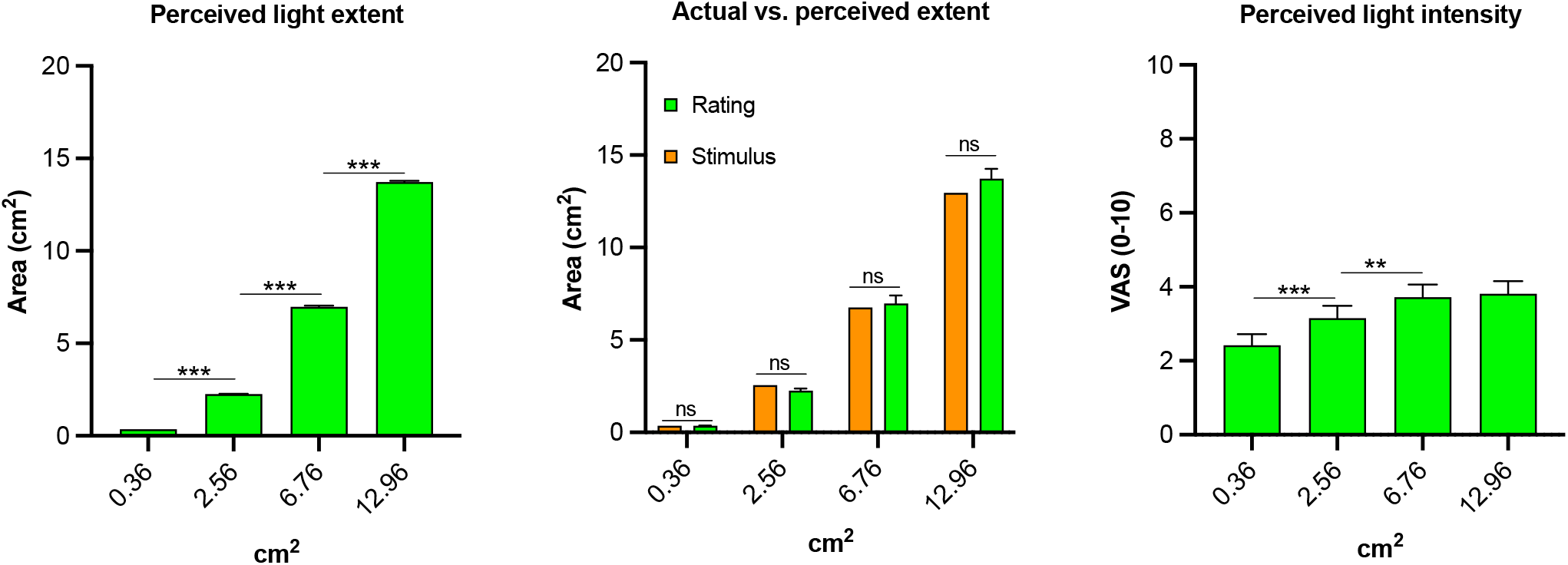
Perceived light extent was remarkably accurate. There is a clear stimulus-response relationship between the actual extent of the displayed object (cm^2^) and the perceived light extent (left panel). Also, participants were very accurate in their light extent ratings: One-sample -Bonferroni corrected- *t* tests performed on perceived light extent against the actual size of light (0.36, 2.56, 6.76, 12.96cm^2^) did not reveal significant differences (middle). Light was generally perceived as more intense when the displayed object (square) was of greater magnitude indicating partial spatial summation of light perception. Significance level: ^*\*\*\**^*p* < 0.001, ^**^*p* < 0.01. Error bars depict standard error of the mean (SEM).

Furthermore, participants were very exact in their size estimations at all light stimulus sizes, with no statistically significant differences between estimate and actual stimulus size: 0.36cm^2^ [*t*_(49)_ = 1.07, *p* = 0.29, mean diff. = 0.03, 95%CI: -0.02 to 0.08], 2.56cm^2^ [*t*_(49)_ = -0.17, *p* = 0.86, mean diff. = 0.04, 95%CI: -0.46 to 0.38], 6.76cm^2^ [*t*_(49)_ = 1.76, *p* = 0.09, mean diff. = 1.32, 95%CI: -0.19 to 2.83], 12.96c,^2^ [*t*_(49)_ = 2.02, *p* = 0.049/ns after Bonferroni, mean diff. = 1.90, 95%CI: 0.01 to 3.78]. Interestingly, greater size of the visual object was to some degree related to perceived light intensity which could suggest involvement of spatial summation: significant effect of “stimulus/size” (F_(3, 147)_ = 30.33, *p* < 0.001, *η*^2^_p_ = 0.38, **Figure 7**). Namely, larger square of light was perceived as more intense following the logarithmic growth: 0.36cm^2^ vs. 2.56cm^2^ [*p* < 0.001, mean diff. -0.73, 95%CI: -1.17 to -0.30], 2.56cm^2^ vs. 6.76cm^2^ [*p* < 0.01, mean diff. -0.57, 95%CI: -1.01 to -0.13], 6.76cm^2^ vs. 12.96cm^2^ [*p* = 1.00, mean diff. -0.09, 95%CI: -0.48 to -0.31].

### 3.6. Psychophysical summary

Steven’s power law was fitted to the data at the subject level and exponents were extracted. In general, pain intensity, unpleasantness and pain extent exhibited exponential increases with an exponent close to m = 4. In, contrast, perceived light intensity was described by exponent lower than 1, and light extent closer to 1, thus approximating a linear relationship **(Figure 8)**.

**Figure 8.**
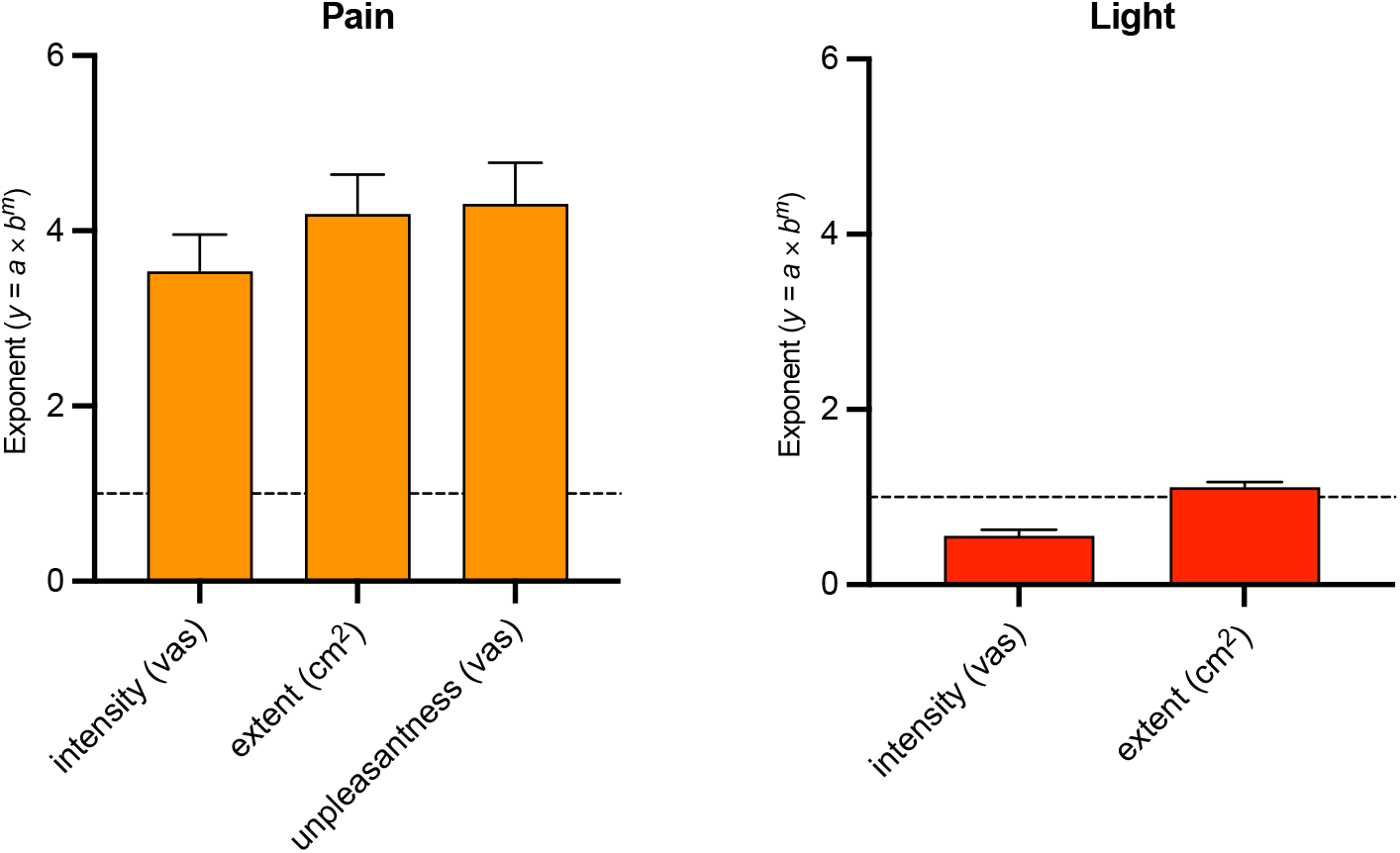
Mean exponents extracted from power functions. The horizontal dashed line represents an exponent = 1, indicating a fully linear relationship between stimulus and perception. Exponents higher than 1 indicate exponential growth, while those lower than 1 suggest logarithmic increase. Perceived pain intensity and unpleasantness exhibited steep exponential growth, consistent with recent findings on larger sample [12]. Similarly, pain extent increased exponentially with temperature (left panel). Conversely, perceived light intensity followed logarithmic increases, aligning with psychophysical literature [41].

### 3.7. Tactile acuity and pain extent

No significant correlations were found between the 2PD thresholds measured at the hand site and the magnitude of pain radiation in response to 10s (*r* = 0.06, *p* = 0.66) as well 5s stimulation (*r* = 0.08, *p* = 0.58). Similar results were noted at the forearm site: 10s stimuli (*r* = 0.01, *p* = 0.96), 5s stimuli (*r* = -0.09, *p* = 0.54).

## 4. DISCUSSION

Noxious thermal stimuli produced pain that extended far beyond the confines of the stimulated skin region, with a 12× larger pain extent than the area of the skin stimulated. This pain radiation effect is present with temperatures higher than 45°C and can be easily evoked at glabrous and non-glabrous skin as well with 5 and 10s stimulation. Within subjects, pain radiation increased as noxious stimulus intensity increased as predicted by population coding mechanisms. However, between-individual differences in perceived pain intensity were only weakly related to the pain radiation, leaving up to 85% of pain radiation accounted by factors other than pain sensitivity. Thus, these spatial attributes of pain may likely involve population coding mechanisms largely distinct from those that encode pain intensity.

Our observations echo early psychophysical observations. The first experimental evidence for the ability of cutaneous pain to expand dates back to 1949 when Kunkle et al. [15] reported in a sample of medical students that following immersion of single digit into cold water some individuals reported pain not only in stimulated digit, but also in the adjacent one. This effect persisted even though the adjacent finger was topically anaesthetized. In the next related investigation Price et al. [31] used graded noxious thermal stimuli to induce pain in 5 healthy participants. No increase in skin temperature was observed 2-3mm from the probe’s edge [31], consistent with evidence that skin cooling mechanisms such as local vasodilation prevent temperature from spreading [18] and isolate the stimulated skin from surrounding areas [11] comparably to laser stimulation [10]. In the Price et al. study [31], approximately 60% of trials (49°C of 5s duration) were associated with radiation of pain. Higher temperatures of 51°C were linked to even greater reports (80% of trials) of pain radiation. Whether the spread of pain generalizes to sensory modalities other than heat and cold remains to be tested. However, referred pain is larger with more intense [29] and longer [9] mechanical stimulation.

### 4.1. Duration of noxious stimulation

More recently, two studies examined pain extent in response to a temporal summation paradigm using noxious heat (or cold) stimuli. The extent of pain increased over repeated stimulus applications [26]. In the subsequent study Naugle et al. [25] found that this effect is positively associated with the age and is more prevalent in older women in particular. To some extent, our results are in line with previous studies with regards to the effect of duration of exposure to heat, however, we did not find any association between pain extent and sex or age. This negative finding should be cautiously interpreted as the study itself was not powered to investigate effects of sex and age. Thus, future studies should investigate these effects prospectively in different age groups.

In the current study, longer noxious heat stimulation led to greater pain radiation. Receptive fields (RFs) of spinal neurons have been shown to expand in response to nociceptive stimulation. Expansion is typically not immediate [13] and takes place within minutes of stimulation with bradykinin [13] or mustard oil [38], although relatively brief (20s) irritation of C fibers can evoke up to 400% increases in RFs of the spinal nociceptive neurons in rats [7]. These increases in RFs sizes may contribute to recruitment of larger populations of spinal cord neurons and could potentially provide a substrate for the radiation of pain.

### 4.2. The effect of stimulation site

Pain extent was markedly different at hairy skin (forearm) vs. glabrous skin (palm). Two peripheral factors could potentially contribute to these differences. First, A-δ fiber innervation differs between hairy and glabrous skin. Hairy skin has both A mechano-heat sensitive Aδ-fibers type I (I AMHs) and type II (II AMHs), while glabrous skin has only AMHs type I. However, the RF sizes of AMH-II vs. AMH-I fibers are not different [43] and therefore could not contribute to differential pain radiation. Second, C-fibers are present in both hairy and glabrous skin and exhibit a distal-proximal gradient of RFs size. Distal areas have smaller RFs sizes than proximal areas [39]. However, in the study by Treede et al. [43] RFs sizes of both A-δ and C-fibers were not different between hairy and glabrous skin. This leaves the impact of RFs sizes of primary afferents (heat-responsive) involvement in pain extent somewhat uncertain. However, peripheral determinants are likely superseded by central mechanisms. The hand is more precisely represented somatotopically within SI compared to the forearm, and has smaller noxious 2PD thresholds and localization errors [20]. This finer spatial tuning in the hand may result in more focused neuron recruitment and lesser radiation of pain.

### 4.3. Accuracy of spatial extent ratings

Control stimuli in the form of visual objects (squares) projected onto the forearm allowed us to quantify measurement error as well as illustrate the absolute accuracy of the extent rating procedure. Results showed that a visual stimulus is rated on average with 0.04cm2 precision, meaning that the error the participant made during extent rating procedure was likely smaller than 1% of the actual size of the object. When visual stimuli were presented with different dimensions from 0.36cm2 to 12.96cm2 - their ratings were accurate: one-sample comparisons did not show significant differences from the size of the target. This nearly exact “matching” is even more surprising considering that the rating procedure was performed after the stimulus was turned off, and therefore required a memory component. Thus, when generalized to pain extent ratings, errors related to the act of making a rating account for a negligible amount of variability of reports of the pain extent, at least in healthy individuals. Future studies should examine this paradigm in individuals with chronic pain.

### 4.4. Pain radiation: Individual differences

Individual differences in intensity of pain only marginally contributed to the variance in pain extent (∼15% of variability). This finding holds substantial significance: First, it suggests that factors linking the experienced pain extent to stimulus extent are distinct from factors linking experienced visual extent to stimulus extent. Second, it indicates that different sources of errors (such as those related to the delivery of the stimulus, reception, appraisal, and manual operation of the trackball) combined contribute to less than 0.01% of the explanation for the variability in pain extent. In summary, one can conclude that the remaining 85% of the variability in pain extent is attributed to factors other than pain intensity, unpleasantness, and error in rating. Thus, the subjective experience of pain extent is largely independent from the subjective experience of pain intensity or unpleasantness. Previous studies have also found virtually no relationship between the magnitude of spatial summation of pain and individual differences in pain intensity [1,34]. The existence of such a substantial unexplained source of information suggests the presence of an independent set of mechanisms responsible for extracting spatial information within the DNS. Indeed, the spatial features of pain are processed via a dorsal stream involving the posterior parietal cortex and dorsolateral prefrontal cortex, while nonspatial features, such as pain intensity, are processed by a ventral stream involving the anterior insular cortex and prefrontal cortex [17,28].

### 4.5. Population coding

The number of neurons recruited by a given noxious stimulus is likely determined by several factors. First, anatomical factors likely define the limits of recruitment. In the spinal cord, such factors would include primary afferent arborization as well as propriospinal interconnections [5,24]. Second, within these limits, the relative contribution of excitatory and inhibitory processes will dynamically define the functional receptive field sizes of individual neurons at a given point in time. Increasing receptive fields sizes would result in increasing recruitment of neurons within the population, while decreasing receptive field sizes would reduce the number of neurons recruited. Psychophysical evidence suggests that inhibition can minimize spatial summation of pain [22] and reduce the recruitment of nociceptive neurons [2]. Consistent with this finding, low noxious stimulus intensities produced a perceived extent of pain that was smaller than the body region stimulated, suggesting that inhibitory processes were dominating. Conversely as noxious stimulus intensities exceeded 45°C the perceived area of pain was substantially larger than the stimulated area, suggesting that excitatory neural processes began to dominate. Consistent with predictions related to the recruitment of spinal cord activity in animal [6] and human [40] models, pain continued to spread as stimulus intensities increased to 49°C.

More widespread neuronal recruitment can not only explain extensive radiation observed in the current study, but also other sensory phenomena such as “filling-in” wherein the DNS interpolates the missing information. In the study by Quevedo & Coghill [34] separation of two nociceptive foci by 10cm contributed to feeling of contiguous pain without actual stimulation. This form of interpolation is possible through increased activation in populations of neurons and can account for clinical phenomena such as widespread pain **(Appendix 1)**.

In summary, sensory features of pain can provide important insights into the nervous system mechanisms giving rise to these experiences. The relative independence of the spread of pain from perceived pain intensity indicates that largely distinct mechanisms are involved in the spatial representation of pain versus perceived pain intensity.

## 5. ACKNOWLEDGEMENTS

The authors would like to thank Wolfgang Loew for his constant help and support with the equipment. Additionally, the authors express their gratitude to Sebastian Elmo Galuszka for his assistance in developing the system to deliver visual stimuli during the study. The authors declare no conflicts of interest.

