## Supplementary materials for "Radiation of pain: Psychophysical evidence for a population coding mechanism in humans"

### Appendix (supplementary materials)

#### S1. Results from General Linear Model performed on pain unpleasantness ratings

| Main or interaction effect | df | *F* | *P* |
|---|---|---|---|
| "site" | 1, 49 | 24.12 | < 0.001 |
| "duration" | 1, 49 | 15.46 | < 0.001 |
| "temperature" | 1.28, 62.88 | 154.42 | < 0.001 |
| "site" × "duration" | 1, 49 | 5.90 | < 0.05 |
| "site" × "temperature" | 1.89, 92.40 | 13.22 | < 0.001 |
| "duration" × "temperature" | 1.87, 91.70 | 14.05 | < 0.001 |
| "site" × "duration" × "temperature" | 2.15, 105.56 | 7.45 | < 0.001 |

Note: The same pattern of results was found in the analysis of pain unpleasantness as in the main analyses conducted on "pain intensity" ratings. All main and interaction effects were statistically significant. Degrees of freedom (df), presented with or without Greenhouse-Geisser correction.

#### S2. Results from General Linear Model performed on light intensity ratings

| Main or interaction effect | df | *F* | *p* |
|---|---|---|---|
| "site" | 1, 41 | 1.39 | 0.25 |
| "duration" | 1, 41 | 0.76 | 0.39 |
| "lux/intensity" | 1.47, 60.19 | 85.54 | < 0.001 |
| "site" × "duration" | 1, 41 | 0.54 | 0.47 |
| "site" × "lux/intensity" | 2.44, 99.97 | 0.59 | 0.59 |
| "duration" × "lux/intensity" | 2.18, 89.25 | 0.67 | 0.53 |
| "site" × "duration" × "lux/intensity" | 2.42, 99.34 | 1.46 | 0.24 |

Note: The table presents detailed results from General Linear Model analysis. Only significant effect of "intensity" was found indicating that participants perceived light differently depending on the level of stimulation (Lux). Some effects are presented following Greenhouse-Geisser correction.

#### S3. Results from General Linear Model performed on light extent ratings

| Main or interaction effect | df | *F* | *p* |
|---|---|---|---|
| "site" | 1, 41 | 0.19 | 0.66 |
| "duration" | 1, 41 | 0.91 | 0.35 |
| "lux/intensity" | 1.36, 55.64 | 11.57 | < 0.001 |
| "site" × "duration" | 1, 41 | 5.59 | < 0.05 |
| "site" × "lux/intensity" | 1.50, 61.45 | 1.74 | 0.19 |
| "duration" × "lux/intensity" | 3, 123 | 1.69 | 0.17 |
| "site" × "duration" × "lux/intensity" | 3, 123 | 1.93 | 0.13 |

Note: The table presents detailed results from General Linear Model analysis. Significant effect of "lux/intensity" was found indicating that participants perceived light extents differently depending on the level of stimulation. Some effects are presented following Greenhouse-Geisser correction.

### S4. Flow-chart and recruitment flow

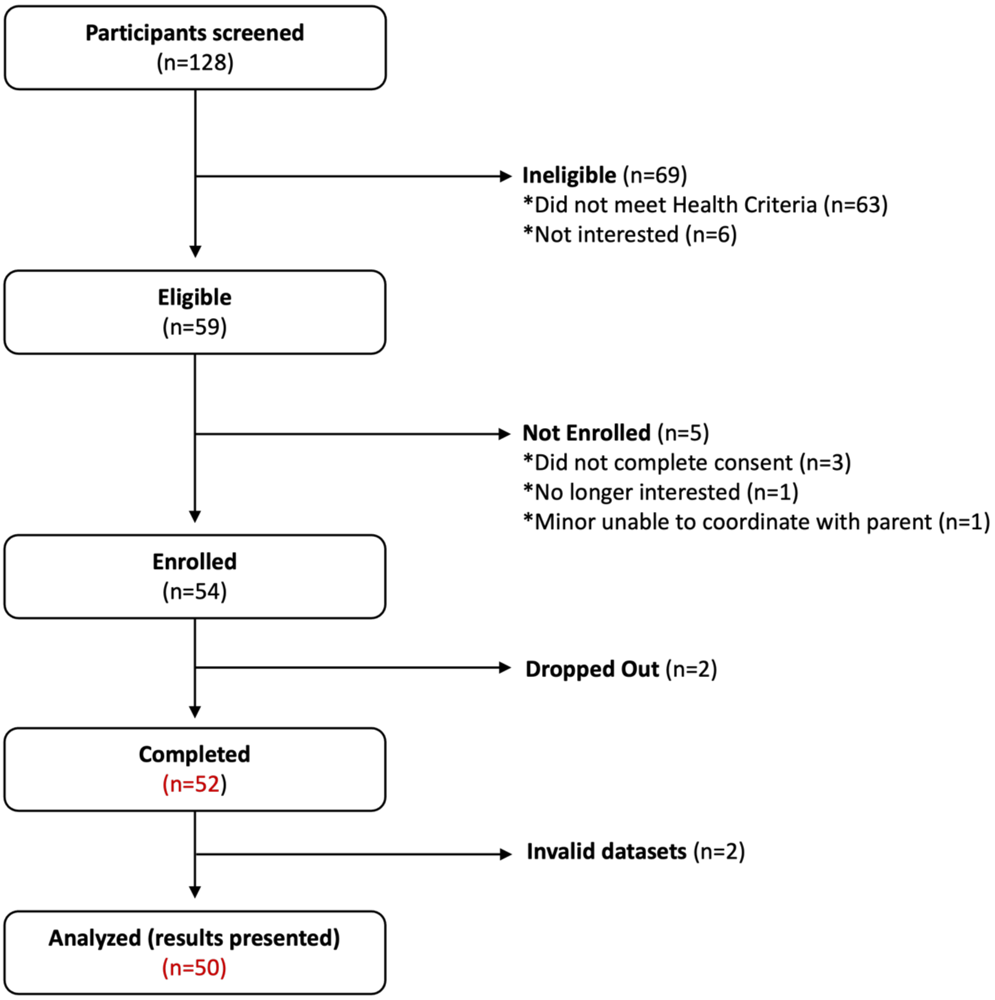


Note: The flow chart has been prepared in line with the CONSORT flow diagram [3].

### S5. Sample size calculation

Sample size was calculated by using a pilot dataset from seven individuals. Similarly to the main experiment participants were exposed to 4 different levels of noxious intensity (43, 45, 47, 49°C) [1]. Each temperature was repeated three times and average outcome was encompassed by power analysis. Power analysis was conducted by simulations performed by means of "Superpower" library [2] implemented in R (run via R studio Version 2023.12.0+369). The code database to reproduce power analysis can be found on GitHub.

### S6. Sample size database

| Site | Variable / condition | meanExtent | sdExtent | Variable / condition | meanExtent | sdExtent |
|---|---|---|---|---|---|---|
| Forearm | 5s 43°C | 0.45 | 0.36 | 10s 43°C | 0.36 | 0.38 |
| | 5s 45°C | 0.55 | 0.49 | 10s 45°C | 0.72 | 0.65 |
| | 5s 47°C | 1.25 | 0.71 | 10s 47°C | 1.74 | 0.76 |
| | 5s 49°C | 2.13 | 1.18 | 10s 49°C | 2.94 | 1.26 |
| hand | 5s 43°C | 0.49 | 0.77 | 10s 43°C | 0.49 | 0.81 |
| | 5s 45°C | 0.57 | 0.73 | 10s 45°C | 0.85 | 0.85 |
| | 5s 47°C | 0.75 | 0.75 | 10s 47°C | 1.40 | 1.20 |
| | 5s 49°C | 1.39 | 0.66 | 10s 49°C | 2.47 | 1.67 |

Note: Data presented here are also stored in the released file on GitHub: /painSize_power_08072022.xlsx. "meanExtent" refers to average pain extent from seven individuals, "sdExtent" refers to standard deviation.

### S7. Manual to access rating task with MacOS compatible version

To run the Python-based rating task, used for the data collection in the current study, follow the steps below. Note that the presented solution was tested on a MacBook Pro (14") with an Apple Silicon M3 chip running macOS Sonoma 14.2.1 (last tested on 2024-03-21):

1. Follow the steps outlined here to install "brew", "python3.10," and all other dependencies.
2. Run this command in your terminal: "source /setHereYourPath/psychopy-on-M1/venv/bin/activate"
3. Note that "venv" is within your cloned object "psychopy-on-M1" (see step #2).
4. Download "painSizeRatingTask_20240425.py" from GitHub.
5. Run it through Python in the terminal: "python /setHereYourPath/painSizeRatingTask_20240425.py"
6. If you want to make changes to the rating task, you can modify the "painSize_instructions_20240425.py" file.
7. The task includes three iterations: i) pain extent rating, ii) intensity, and iii) unpleasantness of pain, and stores ratings within the created directory "/pwd/subjectID".

### S8. Manual to data repository and analyses files

All supporting files related to the current manuscript can be found in the "PAINSIZE" repository on GitHub:

1. Directory "painSizeStats" includes syntax, database in the *.sav format and output of analyses run in the order of presentation in the manuscript
2. Directory "painSizePower" includes R script to run power calculations and a relevant ReadMe file as well as database
3. Directory "painSizeRating" includes "painSizeRatingTask_20240425.py" (see **S7** for details) and modifiable instructions to the task. The task requires Python3 to run.
4. Directory "painSizetMatlab" has all 11 MATLAB *.m files (version 2023b). Note that to run pre-processing of the data and extract a database for relevant statistical analysis one must first initialize the main script "painSize_main.20240305.m" and collect raw data for all 50 individuals. The file may require adjustment of the files' directories, especially directory in which you locally store raw dataset. For further details on how to reproduce analysis from raw level to the data presented on figures follow the "painSize_reproducibility_ReadMe.txt".

### S9. Distribution of the pain extent individual effects across sex and age subgroups.

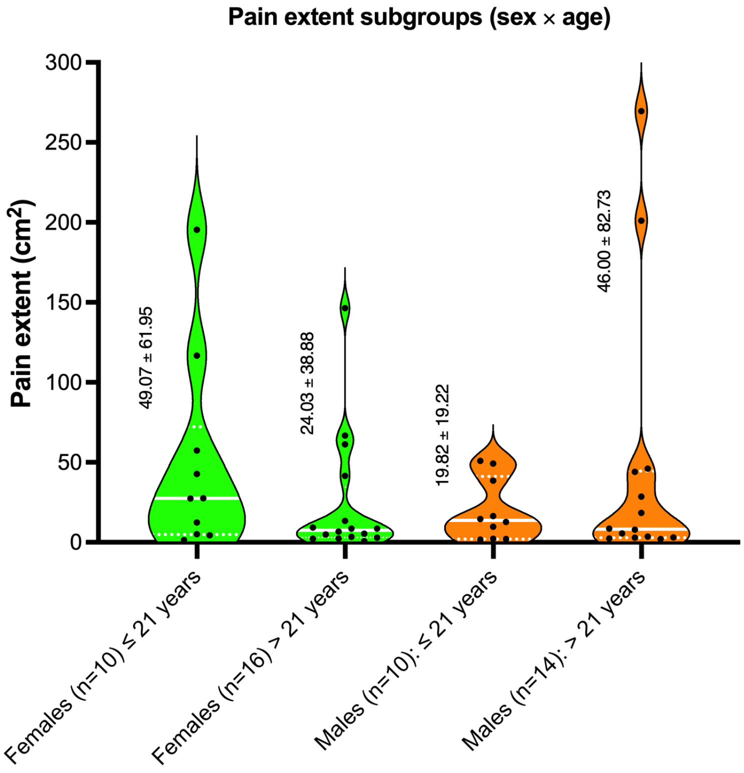


Note: Pain extent across different age subgroup of males and females. Medians are displayed as the white solid lines, while inter-quartile ranges are displayed by the dotted white lines in the violin plot. Standard deviations and mean values are presented on the left side of each violin plot.

### S10. Discussion: clinical implications

The individual differences in the spread of pain evoked by a stimulus of fixed intensity may have substantial clinical implications. Some individuals exhibit modest spread of pain whereas others exhibit extensive spread of pain in a fashion that is largely independent from their perceived pain intensity. This raises the possibility that patients with extensive spreading phenotype may be at greater risk of significant spread of clinical pain. Patients with widespread clinical pain have worse trajectories of recovery than those with more focal pain [5]. Conversely, this spread of pain may also have a protective effect in the event of life-threatening conditions. Patients who experience pain radiating to the arm and face during the heart attack are more likely to seek medical help than those with silent myocardial infraction [4].
